# Multiple parasitoid species enhance top-down control, but parasitoid performance is context-dependent

**DOI:** 10.1101/2021.07.16.452484

**Authors:** Mélanie Thierry, Nicholas A. Pardikes, Miguel G. Ximénez-Embún, Grégoire Proudhom, Jan Hrček

**Affiliations:** University of South Bohemia, Faculty of Science, České Budějovice, Czech Republic; Institute of Entomology, Biology Centre of the Czech Academy of Sciences, České Budějovice, Czech Republic; Georgia State University-Perimeter College, Department of Life and Earth Sciences, 55 North Indian Creek Drive, Clarkston, Georgia 3002

**Keywords:** community composition, community modules, *Drosophila*, interaction modification, multiple predator effects

## Abstract

1. Ecological communities are composed of many species, forming complex networks of interactions. Current environmental changes are altering community composition. We thus need to identify which aspects of species interactions are primarily driven by community structure and which by species identity to predict changes in the functioning of communities. Yet, this partitioning of effects is challenging and thus rarely explored.
2. Here we disentangled the influence of community structure and the identity of co-occurring species on the outcome of consumer-resource interactions using a host-parasitoid system.
3. We used four community modules that are common in host-parasitoid communities to represent community structure (i.e., host-parasitoid, exploitative competition, alternative host, and a combination of both exploitative competition and alternative host). We assembled nine different species combinations per community module in a laboratory experiment using a pool of three *Drosophila* hosts and three larval parasitoid species. To investigate the potential mechanisms at play, we compared host suppression and parasitoid performance across community modules and species assemblages.
4. We found that multiple parasitoid species enhanced host suppression due to sampling effect, weaker interspecific than intraspecific competition between parasitoids, and synergism. However, the effects of community structure on parasitoid performance were species-specific and dependent on the identity of co-occurring species. Consequently, multiple parasitoid species generally strengthen top down-control, but the performance of the parasitoids depends on the identity of either the co-occurring parasitoid species, the alternative host species, or both.
5. Our results highlight the importance of preserving predator diversity for ecosystem functioning, but also show that other effects depend on community composition, and will therefore be likely altered by current environmental changes.

## Introduction

In nature, species interact in a variety of ways, forming complex ecological networks (Fontaine et al., 2011; García-Callejas et al., 2018; Kéfi et al., 2012, 2015; Miele et al., 2019; Pilosof et al., 2017). How species interact depends on the structure of the community, but also on the identity of species in the assemblage (Bográn et al., 2002). Species ranges and phenologies are nowadays quickly shifting in response to environmental changes (Parmesan & Yohe, 2003). Differences in the sensitivity to these changes among species are disrupting historical patterns of interactions and co-occurrences, creating communities with novel species compositions (Alexander et al., 2015). To accurately predict top-down control and forecast the ecological consequences of changes in the biotic environment induced by global changes, we need to understand which aspects of species interactions are primarily driven by community structure and which are driven by species identity.

Trophic (i.e., direct predation) and non-trophic (e.g., competition, facilitation) interactions act in combinations to shape communities (Thierry et al., 2019) and their dynamics (Kawatsu et al., 2021). For example, a predator might switch prey species with the presence of a competing predator (Siddon & Witman, 2004). Most studies of ecological networks are observational (e.g., Jeffs et al., 2021; Tylianakis et al., 2007) and are typically unable to disentangle the potential mechanisms driving species interactions. The strength of top-down control might be driven by a single influential predator species (Letourneau et al., 2009), be enhanced with multiple predator species (i.e., risk enhancement for the prey) if predators show some synergism or degree of niche differentiation (Greenop et al., 2018), or be weakened when multiple predator species are present (i.e., risk reduction for the prey) due to exploitative competition, interference or intraguild predation (Sih et al., 1998). Experimental systems are thus needed to understand the mechanisms structuring networks of interacting species. For this purpose, community modules (i.e., a small number of species interacting in a specified pattern; Holt, 1997, also referred to as “motifs”; Milo et al., 2002) represent a powerful tool to isolate key interactions that structure complex networks. Modules are the building blocks of natural communities (Gilman et al., 2010) and thus allow us to disentangle the structuring mechanisms. Common community modules in food webs include pairwise predator-prey (direct interactions), two prey species sharing a common natural enemy (indirect interactions; i.e., apparent competition or mutualism; hereafter alternative host module), or two predator species attacking the same prey species (either direct or indirect interactions; i.e., exploitative competition, interference, or facilitation; hereafter exploitative competition module). But experimental studies investigating the mechanisms structuring interactions with community modules rarely consider potential variations due to species-specific effects (but see Bográn et al., 2002; Snyder et al., 2006). Thus, it is unclear whether the mechanisms structuring interactions are consistent when looking at community modules of different species compositions (Cusumano et al., 2016).

Experiments manipulating interactions in different community contexts with different species assemblages usually manipulate species assemblage of one trophic level at a time. For instance, Bográn et al. (2002) revealed competitive interactions among predator species in some but not all the predator assemblages studied. However, the study used only one prey species. Snyder et al. (2006) found varying strength in the effect of predator species diversity on aphid suppression depending on the aphid species considered but did not vary species composition in the multiple predator treatment. Woodcock & Heard (2011) showed that predation risk for planthoppers was reduced in systems where predators shared the same habitat domain but did not manipulate the herbivore community. We need studies manipulating species assemblages at both trophic levels to understand how the identity of co-occurring species affects the outcome of consumer-resource interactions.

Here we investigated the mechanisms structuring consumer-resource interactions using a host-parasitoid system. Parasitoids are a diverse group of insects that use other arthropods as a nursery for their offspring, ultimately killing their host to complete development (Godfray, 2004). Parasitoids are important for top-down control in agricultural and natural ecosystems, and are commonly used as biological control agents. Interactions between hosts and parasitoids are easily observed and represent a valuable model system to study how the structure and composition of communities influence species interactions. We used a set of three *Drosophila* species and three of their larval parasitoids from a natural tropical community in Australia (Jeffs et al., 2021). Under laboratory conditions, we reproduced isolated subsets of this community to disentangle the different interactions within host-parasitoid communities. We aimed to uncover general effects of community modules in our *Drosophila*-parasitoid system across different species assemblages and detect any species-specific effects depending on the co-occurring species identity (using nine species assemblages for each of the four common community modules in host-parasitoid networks: host-parasitoid pair, exploitative competition module, alternative host module, and combined exploitative competition and alternative host module). We also explored the mechanisms behind the effects of community structure and composition on host-parasitoid interactions and top-down control on the hosts. Specifically, we tested the following hypotheses: (i) host suppression will be higher in the presence of multiple parasitoid species (i.e., exploitative competition module) due to an increased chance that a more efficient parasitoid species will be present (i.e., sampling effect; Letourneau et al., 2009; Pedersen & Mills, 2004). However, (ii) parasitoid performance might decrease in the presence of multiple parasitoid species due to increased competition in multi-parasitism events (i.e., when multiple parasitoids parasitize the same host individual; Harvey et al., 2013). (iii) Interactions between a focal host-parasitoid pair will change with the presence of an alternative host depending on host choice and host switching behavior. (iv) Combined effects of exploitative competition among parasitoids and alternative host species in the four-species module will differ from three-species modules depending on the identity of the co-occurring species (Bográn et al., 2002; Sentis et al., 2017).

## Materials and methods

### Study system

We used cultures of *Drosophila* species and their associated parasitoids collected from two tropical rainforest locations in North Queensland Australia: Paluma (S18° 59.031’ E146° 14.096’) and Kirrama Range (S18° 12.134’ E145° 53.102’) (< 100 m above sea level) (Jeffs et al., 2021). We established *Drosophila* and parasitoid cultures between 2017 and 2018, identified them using morphology and DNA barcoding, and shipped them to the Czech Republic under permit no. PWS2016-AU-002018 from Australian Government, Department of the Environment. We maintained all cultures at 23°C on a 12:12 hour light and dark cycle at the Biology Centre, Czech Academy of Sciences. We used three host species (*Drosophila birchii, D. simulans*, and *D. pallidifrons*), and three larval parasitoid species *Asobara sp*. (Braconidae: Alysiinae; strain KHB, reference voucher no. USNMENT01557097, reference sequence BOLD process ID:DROP043-21), *Leptopilina sp*. (Figitidae: Eucolinae; strain 111F, reference voucher no. USNMENT01557117, reference sequence BOLD process ID:DROP053-21), and *Ganaspis sp*. (Figitidae: Eucolinae; strain 84BC, reference voucher no. USNMENT01557102 and USNMENT01557297, reference sequence BOLD process ID:DROP164-21) (for more details on the parasitoid strains used see Lue et al., 2021). We kept *Drosophila* isofemale lines on standard *Drosophila* medium (corn flour, yeast, sugar, agar, and methyl-4-hydroxybenzoate) for approximately 45 to 70 non-overlapping generations. We combined four to seven lines from each host species into two population cages of mass-bred lines per species to revive genetic variation before the start of the experiment. We used single parasitoid isofemale lines that we maintained for approximately 25 to 40 non-overlapping generations before beginning the experiment by providing them with two-day-old larvae of *Drosophila melanogaster* every week. This host species was not used in the experiment, thus avoiding potential bias due to maternal effects.

### Experimental design

To investigate the effects of community structure and species composition on host-parasitoid interactions, we used four community modules (Figure 1), each represented by nine different species assemblages, from the pool of three host species and three parasitoid species (six host species assemblages by six parasitoid species assemblages for a total of 36 species assemblages). We replicated each unique treatment six times. Each replicate was represented by a set of two vials in one box, for a total of 216 boxes (N = 432 vials) [N.B.; one replicate with *D. birchii, D. simulans*, and two *Ganaspis sp*. was unsuccessfully set up and therefore removed from the analyses]. We used either conspecific (Figures 1a and c) or heterospecific (Figures 1b and d) parasitoids. The two vials contained *Drosophila* larvae from the same host species (Figures 1a and b) or different host species (Figures 1c and d). We kept host and parasitoid densities constant across treatments (i.e., substitutive design; e.g., Snyder et al., 2008) to focus on the consequence of changes in the diversity of parasitoid and host species that may occur with global change. We did not investigate the effects of changes in parasitoid and host density within communities here. We simultaneously performed controls for each host species without parasitoids to obtain the baseline for host survival without parasitism (replicated eight times each, N = 24 vials).

**Figure 1.**
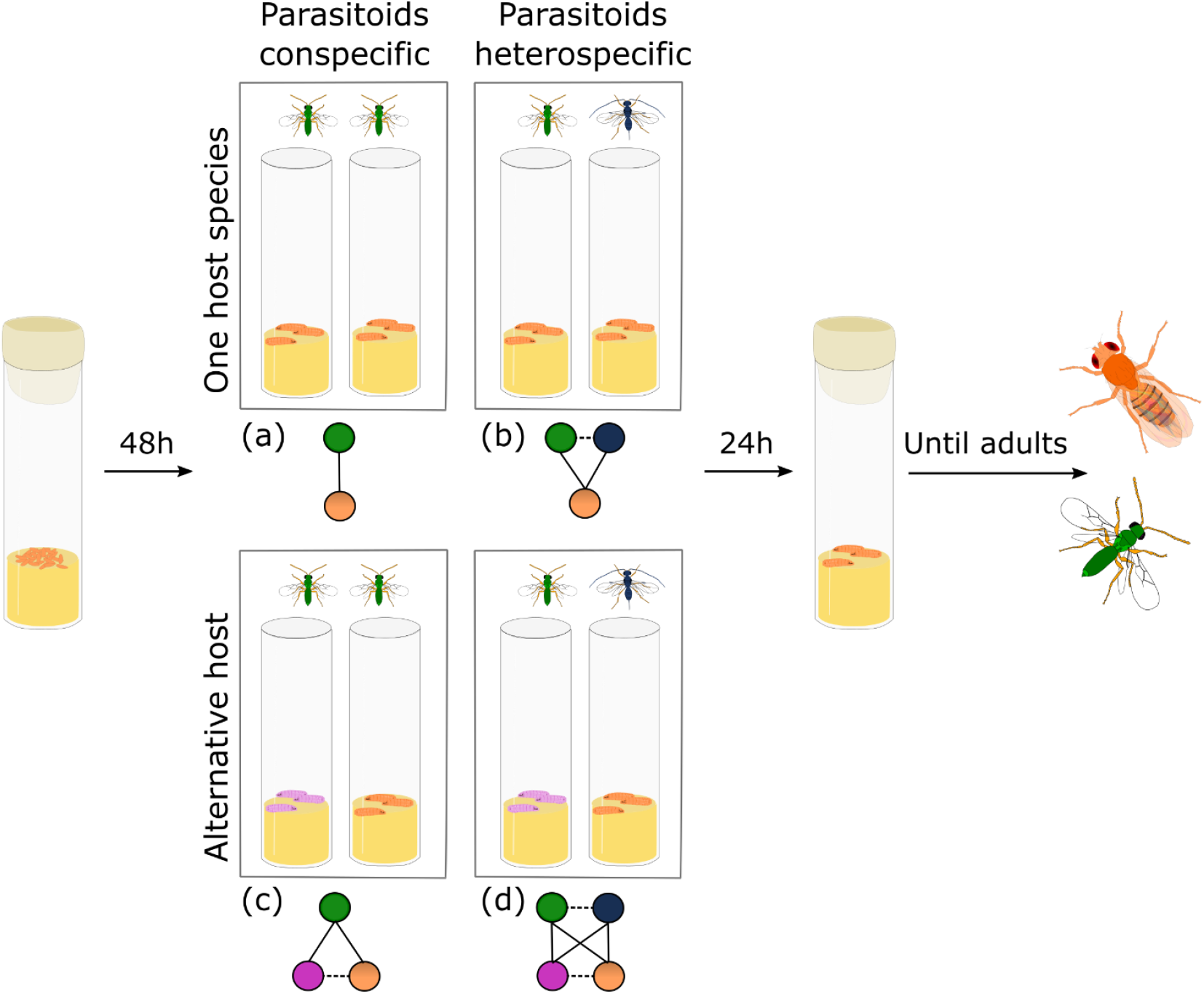
Schematic representation of the experimental treatments with the potential direct and indirect interactions in each community module. Orange and pink nodes and larvae represent different *Drosophila* host species, and green and blue nodes and wasps represent different parasitoid species, assembled in a fully factorial design in four different community modules represented schematically below their corresponding experimental box: a) host-parasitoid pair (one host and one parasitoid species), b) exploitative competition module (one host and two parasitoid species), c) alternative host module (two host and one parasitoid species) and, d) both exploitative competition and alternative host module (two host and two parasitoid species). In the community module schemas, solid lines represent trophic interactions, and dashed lines represent non-trophic interactions (in b) either exploitative competition, interference, or facilitation between parasitoids, c) indirect interactions between hosts, and d) potential for all the above). Direct interaction between host species were not allowed. See Thierry et al., (2019) for a detailed description of each interaction type

We placed twenty-five eggs of each host species in a single 90 mm high and 28 mm diameter glass vial with 10mL of food media to initiate the experiment. We adapted an egg-wash protocol from Nouhaud et al. (2018) to collect *Drosophila* eggs. The day before the egg-washed protocol was conducted, we introduced two Petri dishes with egg-laying medium (agar gel topped with yeast paste) in each population cage for flies to lay eggs overnight. We then transferred eggs to experimental vials.

After 48 hours, we placed two vials with *Drosophila* second instar larvae (initially eggs) in a hermetically sealed plastic box (15 × 11 × 19 cm) with four 3-to-5-days-old parasitoids (1:1 sex ratio). We removed the parasitoids twenty-four hours later. We removed vials from the boxes and plugged them for rearing (Supporting Figure S1). We checked vials daily for adult emergences until the last emergence (up to 41 days for the species with the longest developmental time). We waited five consecutive days without any emergence to stop collecting and thus avoided confounding counts with a second generation. We collected, identified, and sexed all emerges and stored them in 95% ethanol.

We collected a total of 11,400 host eggs across 456 experimental vials. 7,494 eggs (66%) successfully developed into adults (3,702 hosts and 3,792 parasitoids). The remaining 34% either died naturally, or the host was suppressed without successful development of the parasitoid.

### Data analysis

We characterized the outcome of host-parasitoid interactions by a combination of Degree of Infestation (DI; i.e., host suppression) and Successful Parasitism rate (SP; i.e., parasitoid performance). The Degree of infestation (*DI*) and Successful parasitism rate (*SP*) were measured as:

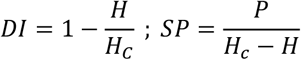

where *H* is the number of adult hosts emerging from the experiment vial, *H*_*C*_ is the mean number of adult hosts emerging from the controls without parasitoids, and *P* is the number of parasitoid adults emerging from the experimental vial (Boulétreau & Wajnberg, 1986; Carton & Kitano, 1981). *DI* was set to zero if the number of hosts emerging from the treatment was greater than the controls. If no parasitoid emerged or the number of hosts suppressed was estimated to be zero, *SP* was set to zero. If the number of parasitoids that emerged was greater than the estimated number of hosts suppressed, *SP* was assigned to one. For treatments with parasitoid conspecifics, we assumed that each of the two parasitoid individuals was attacking the hosts equally; therefore, the number of parasitoid adults emerging was divided by two to calculate individual successful parasitism rate.

We analyzed the data with generalized linear mixed-effects models (GLMMs) and verified model assumptions with the *DHARMa* package (Hartig, 2019). To correct for overdispersion of the residuals, data were modeled using a beta-binomial error distribution and a logit link function using the *glmmTMB* function from the *TMB* package (Lüdecke et al., 2019). We compared the AIC of models with or without zero inflation. We chose the models with the smaller AIC, which were the zero-inflated models. Three model types were used to investigate the general effects of community modules, species-specific responses, and effects of community composition. (i) “Community module models” used two explanatory variables and their two-way interaction to account for the fully-factorial design of the experiment that resulted in four community modules (exploitative competition treatment with two levels: presence or not of a parasitoid heterospecific, and alternative host treatment with two levels: presence or not of an alternative host species). Box ID (214 levels) was included as a random factor to remove the variation between the different species assemblages and thus extract general effects of community modules. Host species (three levels) for DI and host-parasitoid pairs (nine levels) for SP were also included as random factors to remove the variation between different species. (ii) “Species-specific community module models” used the same explanatory variables and Box ID as a random factor as previously described, but host species and host-parasitoid pairs were included as fixed factors to test if effects varied depending on the focal species. All three and two-way interactions between treatments, host species, and host-parasitoid pairs were tested and kept in our models if found to be significant based on backward model selection using Likelihood-ratio tests. Models for SP were also run for each host-parasitoid pair separately to quantify differences in the sign and magnitude of the effects of community structure on pairwise interaction depending on the focal species. (iii) “Community composition models” used species assemblages rather than community modules as explanatory variables (host species assemblage: six levels, and parasitoid species assemblage: six levels). The two-way interaction between host and parasitoid assemblages was always kept in the models to account for the fully-factorial design of the experiment. Models for DI were run for each host species, and models for SP were run for each host-parasitoid pair separately. Blocks (six levels) were included in all models as a random effect. The significance of the effects was tested using Wald type III analysis of deviance with Likelihood-ratio tests. Factor levels of community modules and species assemblages were compared to the reference module and species assemblages of the host-parasitoid pair in isolation by Tukey’s HSD *post hoc* comparisons of all means, using the *emmeans* package (Lenth, 2018).

To further investigate emergent effects of parasitoid diversity on host suppression, we compared the observed and expected outcomes on host suppression when multiple parasitoid species were present, following the framework proposed by Schmitz (2007). Observed values smaller than estimated translate to risk enhancement for the host with multiple parasitoid species, while observed values bigger than estimated reflect risk reduction. Detailed methods are presented in Supporting Text S1. All analyses were performed using R 4.0.2 (Team, 2017).

## Results

### Effects of community structure on host suppression

The presence of multiple parasitoid species in the module significantly increased the probability of hosts being parasitized (DI) (community module model: χ2_(1)_ = 7.08, P = 0.008; Figure 2a; Supporting Table S1). Moreover, effects of multiple parasitoid species on host suppression were greater than expected in four host-parasitoid cases out of nine (Supporting Figure S2). However, DI did not significantly change with the presence of an alternative host species (χ2_(1)_ = 0.56, P = 0.452; Odd Ratio (OR) alternative host module/pairwise interaction = 0.96, P = 0.984), and the two-way interaction between alternative hosts and exploitative competition treatments had no significant effect on DI (community module model: χ2_(1)_ = 0.22, P = 0.638; Figure 2a; Supporting Table S1).

**Figure 2.**
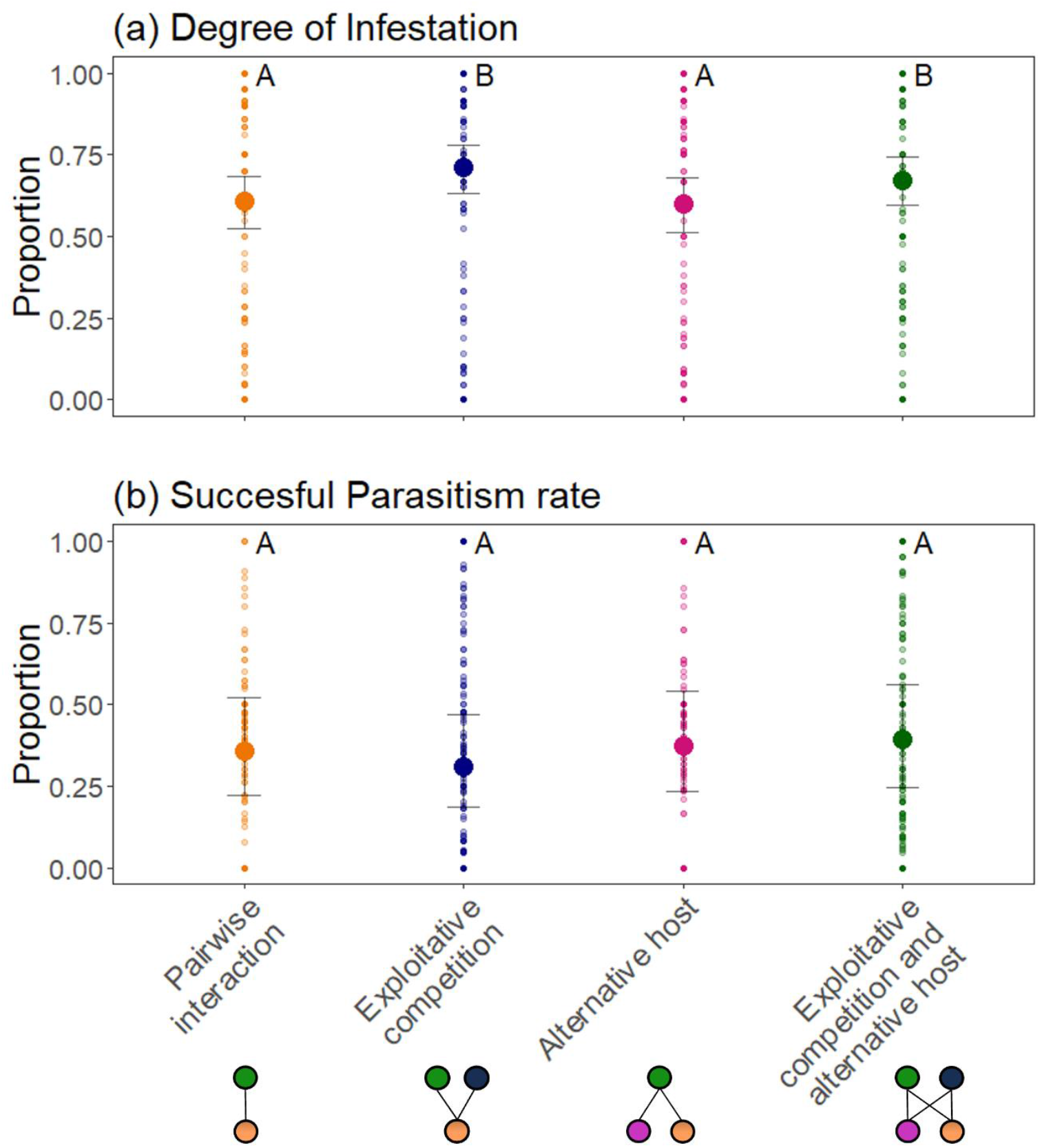
Effects of community structure (represented by the community module bellow each treatment) on (a) degree of infestation (N = 430) and on (b) successful parasitism rate (N = 646). Different capital letters denote significant differences between community structure from the community module models. The small points represent the observed values, and the large points represent the predicted values with their 95% confidence intervals

### Effects of host species and community composition on host suppression

Host DI did not differ significantly across host species (species-specific community module model: χ2_(2)_ = 0.07, P = 0.965; Supporting Table S2). The directionality of the effect of parasitoid diversity did not vary depending on species assemblages (community composition models; Supporting Figure S3 and Table S3).

### Effects of community structure on parasitoid performance

Community modules had no general effects on successful parasitism rates (SP) (community module model; Figure 2b; Supporting Table S4), but the effects significantly varied across host-parasitoid pairs (species-specific community module model; three-way interaction: χ2(8) = 36.81, P < 0.0001, Figure 3, Supporting Table S5). The interaction between exploitative competition and alternative host treatments significantly affected SP for one out of the nine host-parasitoid pairs (*Ganaspis sp*. on *D. simulans*). Successful parasitism rate of two other host-parasitoid pairs significantly decreased with exploitative competition between parasitoid species (*Ganaspis sp*. On *D. birchii* and *D. pallidifrons*), while SP significantly increased with exploitative competition between parasitoid species for *Asobara sp*. on *D. simulans*. Successful parasitism rate for the rest of the host-parasitoid pairs did not significantly change between community modules compared to the host-parasitoid pair in isolation (Figure 3).

**Figure 3.**
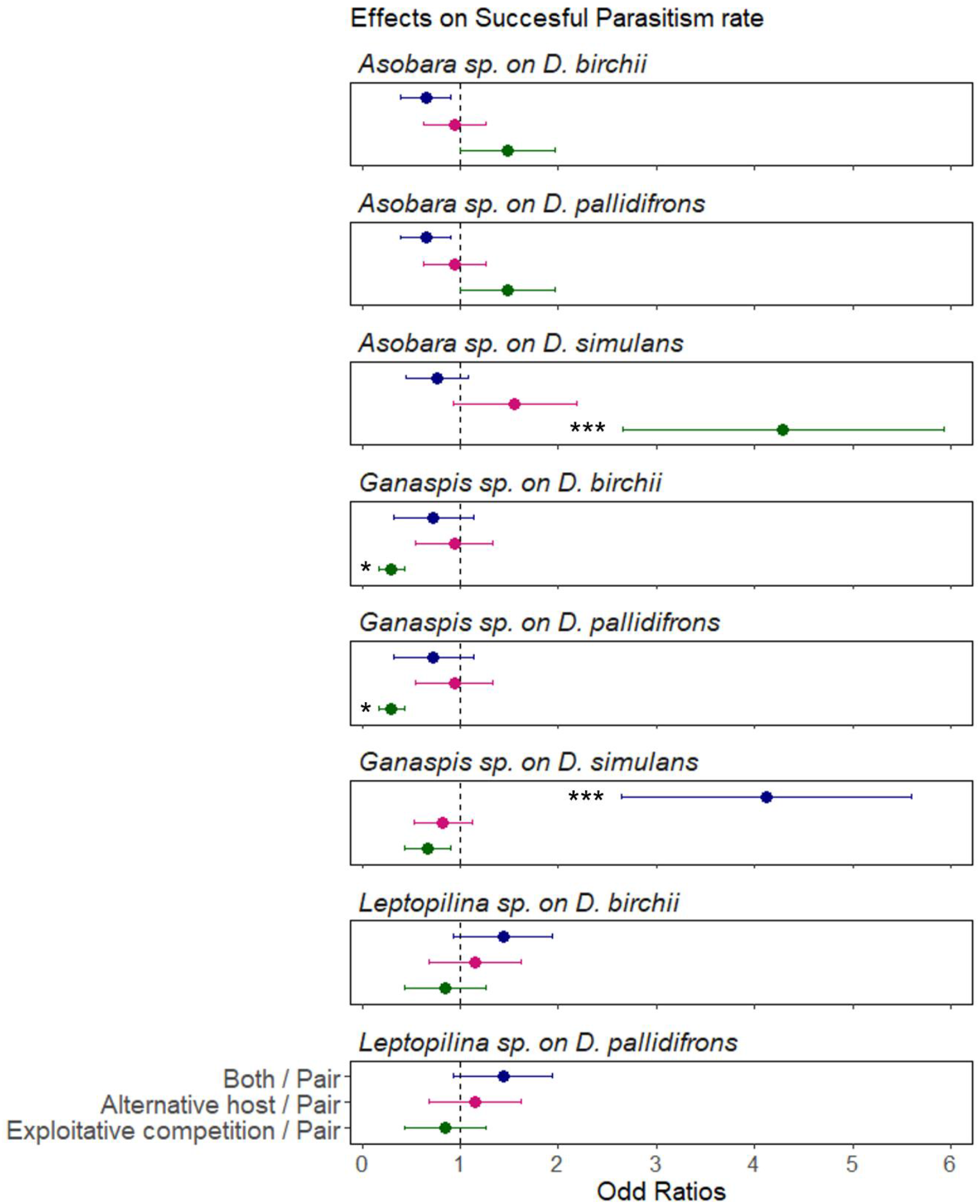
Species-specific effects of community structure on successful parasitism rates. Odds ratios represent effects of each community module (exploitative competition, alternative host, and both exploitative competition and alternative host) on successful parasitism rates compared to the host-parasitoid pair in isolation for each pair (host abbreviations: b: *D. birchii*, p: *D. pallidifrons*, s: *D. simulans*, and parasitoid abbreviations: A: *Asobara sp*., L: *Leptopilina sp*., G: *Ganaspis sp*.). Odds Ratios superior or inferior to 1 translate an increased or a decreased probability of successful parasitism, respectively. Results come from the species-specific community module models run for each host-parasitoid pair separately. Odds Ratios are presented with their standard error and the significance of the effect: (***) P < 0.001, (**) P < 0.01, (*) P < 0.05, (ns) P > 0.05.

### Effects of community composition on parasitoid performance

Effects of an alternative host and a parasitoid competitor on parasitoid performance varied depending on co-occurring species identity, both in magnitude and direction of their response (community composition models; Table 1 and Figure 4). The interaction between host and parasitoid species assemblages had a significant effect on SP for four out of the nine host-parasitoid pairs: *Asobara sp*. on *D. simulans, Leptopilina sp*. on *D. birchii*, and *Ganaspis sp*. on *D. birchii* and on *D. simulans*. Effects of species assemblages on SP for each host-parasitoid pair are presented in Supporting Text S3.

**Table 1.** Effects of community composition on the probability of successful parasitism for each host-parasitoid pair. Effects are shown by the summary of Likelihood-ratio chi-square tests on the community composition models with the effects of host and parasitoid species assemblages (three levels each) (parasitoid abbreviations: A: *Asobara sp*., L: *Leptopilina sp*., G: *Ganaspis sp*. and host abbreviations: b: *D. birchii*, p: *D. pallidifrons*, s: *D. simulans*) (N = 72 for A-b, L-b, A-p, L-p, A-s, L-s, G-s and N = 71 for G-b, G-p). For A-p and L-s models contain only host species assemblage as a fixed effect due to convergence problem with the full model. Degrees or freedom (Df) are given for each factor and for the residuals. χ2 values are presented with the significance of the effect: (***) P < 0.001, (**) P < 0.01, (*) P < 0.05, (ns) P > 0.05.

| Effects | Df | A-b |  | A-p |  | A-s |  | L-b |  | L-p |  | L-s |  | G-b |  | G-p |  | G-s |  |
| --- | --- | --- | --- | --- | --- | --- | --- | --- | --- | --- | --- | --- | --- | --- | --- | --- | --- | --- | --- |
| Host sp. | 2 | 11.14 | ** | 1.12 | (ns) | 15.56 | *** | 4.80 | (ns) | 4.83 | (ns) | 10.08 | ** | 34.14 | *** | 2.23 | (ns) | 19.71 | *** |
| Parasitoid sp. | 2 | 6.63 | * | - |  | 0.11 | (ns) | 38.32 | *** | 4.36 | (ns) | - |  | 2.57 | (ns) | 1.73 | (ns) | 13.81 | ** |
| Host x para | 4 | 7.80 | (ns) | - |  | 24.19 | *** | 40.57 | *** | 7.81 | (ns) | - |  | 25.22 | *** | 9.01 | (ns) | 20.51 | *** |
| Df residuals |  | 60 |  | 66 |  | 60 |  | 60 |  | 60 |  | 66 |  | 59 |  | 59 |  | 60 |  |

**Figure 4.**
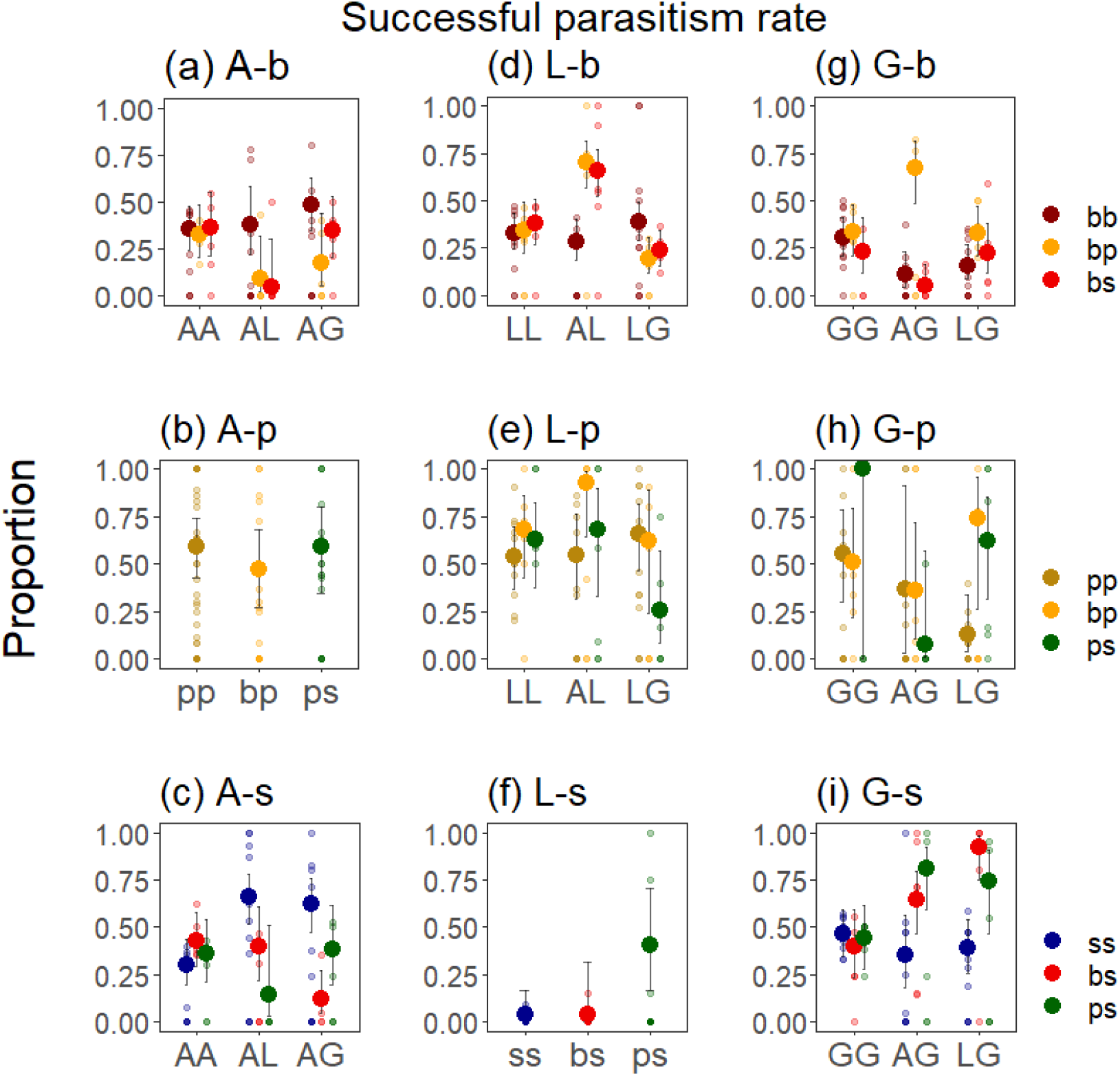
Effects of community composition (identity of the alternative host and the parasitoid heterospecific) on the successful parasitism rate of each parasitoid species on each host species [columns are parasitoid species (parasitoid abbreviations: A: *Asobara sp*., L: *Leptopilina sp*., G: *Ganaspis sp*.) and rows are host species (host abbreviations: b: *D. birchii*, p: *D. pallidifrons*, s: *D. simulans*)] (N = 72 for A-b, L-b, A-p, L-p, A-s, L-s, G-s and N = 71 for G-b, G-p). Host assemblages are represented by the different colors, and parasitoid assemblages are on the x axis. For SP A-p and SP L-s, only effect of host assemblages was analyses due to convergence problem with the full model, and are represented for all parasitoid assemblages combined. The small points represent the observed values, and the large points represent the predicted values with their 95% confidence intervals from the community composition models

## Discussion

Our results confirm some general effects of community structure on consumer-resource interactions over several species combinations and reveal important species-specific effects linked to the identity of species composing the community. Specifically: (i) the presence of multiple parasitoid species consistently increased host suppression due to a combination of mechanisms discussed below. On the contrary, (ii) the presence of an alternative host had no general effect on host suppression but increased or decreased successful parasitism rate depending on the host-parasitoid pairs and co-occurring species identity. These species-specific effects on parasitoid performance have important implications for biological control programs that rarely take into account the impact of co-occurring species on biological control agent efficiency to suppress pests. Moreover, species diversity decrease due to ongoing environmental changes is likely to lead to a decrease in top-down control, and thus changes in the balance between trophic levels in nature.

### Positive effects of consumer diversity on top-down control

The presence of multiple parasitoid species generally increased host suppression (community module model). We found a greater top-down control than expected with multiple parasitoid species for four out of the nine species combinations, suggesting weaker interference between parasitoids heterospecific than between conspecific parasitoids, or synergism between parasitoid species (Snyder et al., 2008; Snyder & Tylianakis, 2012). Weak predator interference can be obtained if predators show some degree of resource partitioning (Pedersen & Mills, 2004), which has been reported in several experimental studies (reviewed in Letourneau et al., 2009). Resource partitioning can be more important than diversity *per se* to explain an increase in top-down control with multiple predator species (Finke & Snyder, 2008). Synergistic effects could also explain a greater top-down control than expected with multiple parasitoid species. Indeed, hosts can overcome a single parasitism event with their immune response but are more likely to die when multi-parasitized (Thierry et al., 2021). The positive effects of parasitoid diversity on host suppression were also driven by the presence of the most efficient parasitoid species for the focal host (e.g., presence of *Ganaspis sp*. for *D. simulans*: Supporting Figure S3c), which could explain why there were no significant differences between observed and expected effect magnitude of multiple parasitoid species in some cases. These results match the sampling effect model suggesting that an increase in top-down control with increased consumer diversity is explained because of an increased probability that a superior natural enemy species will be present in the community (Myers et al., 1989). Here, both the sampling effect, resource partitioning, and synergism between parasitoids seem to be in play to explain the increase of host suppression with an increase in parasitoid diversity. Our results demonstrating the enhancement of top-down control with multiple parasitoid species across several species assemblages, in addition to the results from previous studies (reviewed in Letourneau et al., 2009), highlight the importance of preserving predator diversity for ecosystem functioning.

### No indirect interactions among prey species

Contrary to the presence of an additional parasitoid species, we did not detect any effect of an additional host species on host suppression. This suggests that, in situation with equal host abundances, an alternative host species did not have any effect on host-parasitoid interactions in terms of host switching behaviors. There are other possible effects of an alternative host species on host-parasitoid interactions (i.e., density- and trait-mediated effects; (Werner & Peacor, 2003) that were not tested in our experiment. Yet, they may (Fleury et al., 2004; McPeek, 2019; Morris et al., 2001; van Veen et al., 2005) or may not (Jones et al., 2009; Kaartinen & Roslin, 2013) be important.

### Importance of community composition for consumer-resource interactions

We based our study on a limited set of interacting species, yet even the relatively small number of species used in our experiment allowed us to uncover species-specific responses within a given community module. Community modules have been extensively used as a tool to study the mechanisms structuring and stabilizing complex natural communities (Bascompte & Melián, 2005; Rip et al., 2010), yet the effects of species identity in such studies are often ignored. Our results highlight that variation in directionality and magnitude of the effects of a particular community module on host-parasitoid interactions depend on the species assemblage considered. This is particularly relevant to predicting natural top-down control and the consequences of climate change driven shifts in community composition for ecosystem functioning. Potential effects of co-occurring species (alternative prey species and/or other predators) should be considered when developing biological control strategies, especially when choosing the biological control agent in open agricultural systems. For example, the presence of non-target species might weaken the biological control agent-pest interaction (Wajnberg et al., 2001). Even with no direct effects of the presence of a non-target species, the presence of a natural enemy to the alternative prey species could decrease the biological control agent performance and prevent its establishment in the community. However, the differences between the number of hosts suppressed and the outcome for the parasitoid populations depending on the biotic context might be a unique feature of host-parasitoid systems. It remains to be determined if the identity of co-occurring species is as important in prey-predator systems.

Successful parasitism rate increased in modules with a parasitoid competitor compared to the pairwise interaction in six species combinations, suggesting that some parasitoid species benefit from the presence of a heterospecific. According to a recent review on interspecific interactions among parasitoids (Cusumano et al., 2016), and the best of our knowledge, only one study showed facilitation between two parasitoid species (Poelman et al., 2014). Our case seems different because successful parasitism rates did not increase for both parties, implying competitive interaction between the parasitoids. Parasitoids can compete both as adults for oviposition (i.e., extrinsic competition or interference; Ode et al., 2022) and as larvae within a host (i.e., intrinsic competition; Harvey et al., 2013). Extrinsic competition has adverse effects on parasitoid attack rates, linked to search efficiency and handling time, leading to a potential decrease in host mortality (Xu et al., 2016). In our study, modules with the pairwise interaction in isolation had two parasitoid conspecifics (i.e., substitutive design). Our results suggest that in these six cases, where parasitoid performance increased with the presence of a competitor, interspecific extrinsic competition between parasitoids was weaker than intraspecific extrinsic competition. On the other hand, intrinsic competition results from a super-or multi-parasitism event when two parasitoids (conspecifics or heterospecific, respectively) parasitize the same host individual. It is usually detrimental for the host survival and would explain an increase in host suppression with multiple parasitoid species despite a decrease in parasitoid performance in some cases (i.e., unsuccessful parasitism but with host suppression; e.g., Thierry et al., 2021).

Our contrasting results on successful parasitism rate depending on the host-parasitoid pair and the other parasitoid species present in the community are likely due to differences in traits (e.g., the hosts’ immune response and oviposition behavior and virulence of the parasitoids; Carton et al., 2008). For example, *D. simulans* often encapsulates parasitoid eggs, while the encapsulation rates of *D. birchii* are low in our system. *Asobara sp*. rarely lays more than one egg in the same host individual but develops fast to avoid encapsulation, while super-parasitism is common for *Ganaspis sp*. and could be a strategy of this parasitoid to overcome host immune response. The different trait combinations and trade-offs across host and parasitoid species are likely an important mechanism driving species interactions and co-occurrences in natural communities (Wong et al., 2019).

## Acknowledgements

We thank Anna Mácová, Andrea Weberova, Petr Lavička, and Joel Brown for their help to set up the experiment, Chia-Hua Lue for fruitful discussions on the results. We also thank Owen Lewis, Ben Woodcock and anonymous reviewers for comments that improved the manuscript. Tereza Holicová made the drawings used for Figure 1. We acknowledge funding support from the Czech Science Foundation grant no. 20-30690S.

## Conflict of Interest

The authors declare no competing interests.

## Authors’ contributions

MT conceived the project; NP and JH contributed to the experimental design; MT, NP, MG, and GP collected the data; MT analyzed the data and led the writing of the manuscript. All authors contributed critically to the drafts and gave final approval for publication.

## Data Availability Statement

All raw data and R scripts used for this study are available from the Zenodo database: 10.5281/zenodo.6122978.

## Notes

https://zenodo.org/record/6122978#.Yg5m3ejMJ9A

### Competing Interest Statement

The authors have declared no competing interest.

