## Supplement Materials for "Multiple parasitoid species enhance top-down control, but parasitoid performance is context-dependent"

In *Journal of Animal Ecology*

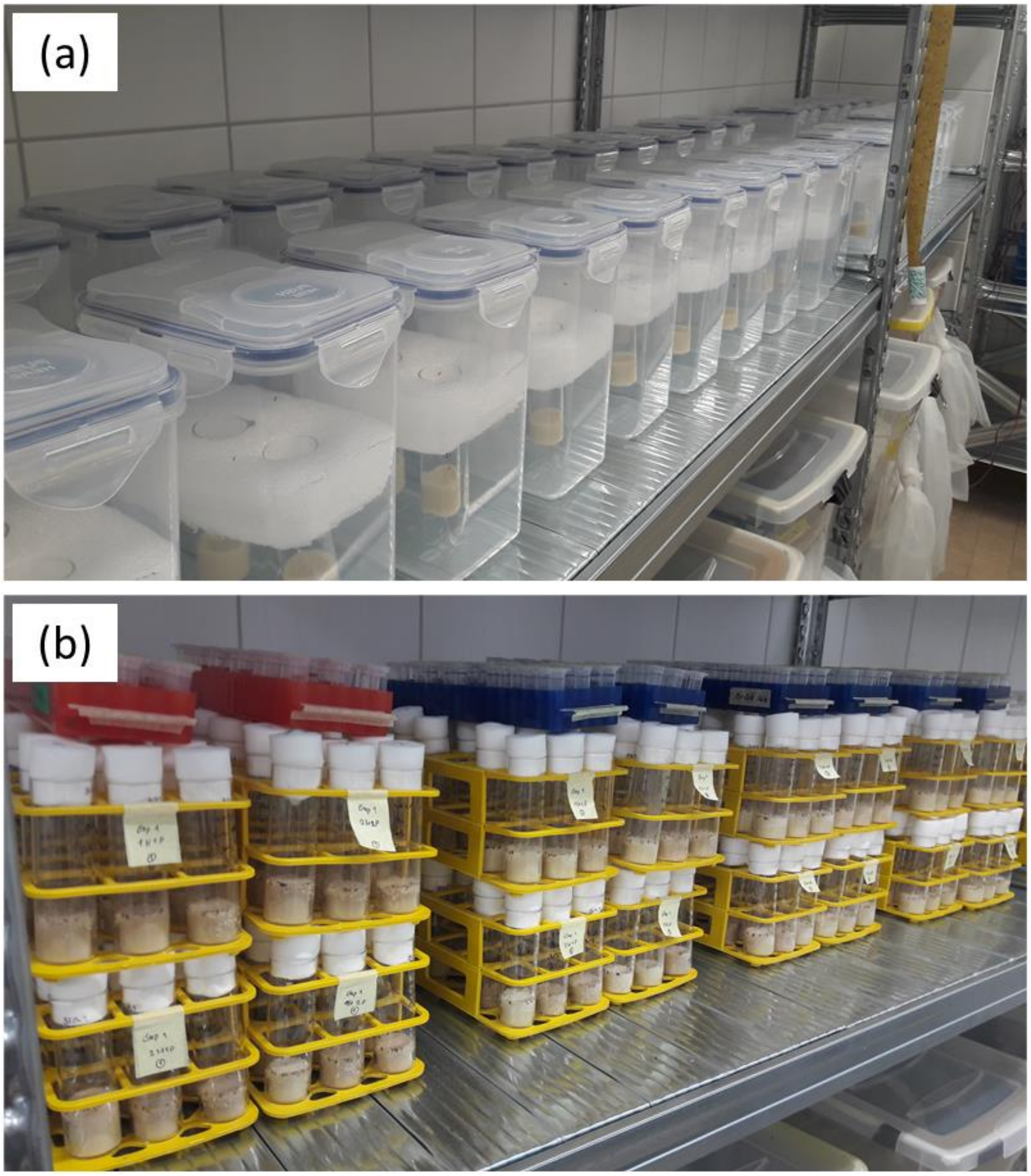

**Figure S1.** *Experimental set up.* (a) Boxes contained two vials with 25 two-days-old Drosophila larvae. Four three-to-five days old parasitoids (1:1 sex ratio) were placed in each box for 24 hours. The experiment counts a total of 216 boxes. (b) After 24 hours, parasitoids were removed, and vials were plugged for rearing. Emerges were collected daily and kept in 95% ethanol.

**Supporting Text S1:** *Emergent effects of parasitoid diversity on host suppression*

Emergent effects of parasitoid diversity on host suppression were analyzed using the approach presented by Schmitz (2007). The observed effect magnitude of multiple parasitoids was calculated as the log ratio of host survival:

$$R_{ij} = \ln(HS_{ij})$$

where $HS_{ij}$ is the observed host survival rate (number of hosts at the end of the experiment with two parasitoids *i* and *j*, either conspecifics or heterospecifics, over number of hosts from the controls). We tested the condition for substitutability:

$$\frac{R_{ii}}{P_i} + \frac{R_{jj}}{P_j} = \frac{4 * R_{ij}}{P_i + P_j}$$

where $R_{ij}$ is the observed effect magnitude of multiple parasitoids *i* and *j*, and $P_i$ is the number of parasitoid *i*. Because there were no significant differences in DI between host treatments (i.e., presence or not of an alternative host species; P > 0.5), we combined data from the different host species combinations (n = 9). We calculated the values for each heterospecific parasitoids assemblage (n = 3), each host species (n = 3), and each replicate (n = 6). We tested significant differences the left- and right-hand sides of the equation for the condition for substitutability with matched-pairs *t* tests. Substitutability can be inferred if the test statistic is not significant.

Effect of multiple parasitoid species on host suppression was greater than expected for *D. birchii* with one out of the three parasitoid combinations (Figure S2a), for *D. pallidifrons* with two out of the three parasitoid combinations (Figure S2b), and for *D. simulans* with one out of the three parasitoid combinations (Figure S2c). For the rest, observed and estimated effects of multiple parasitoid species were not significantly different (Figure S2), indicating substitutability in these five cases.

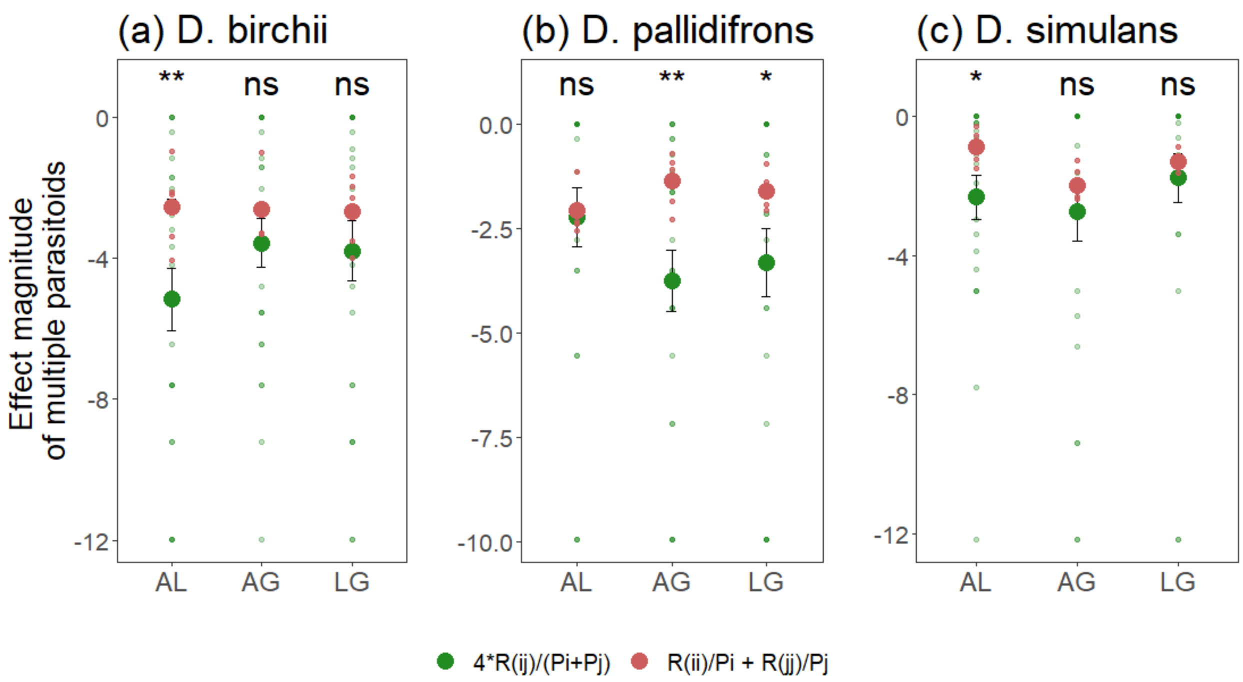

**Figure S2**. *Effect magnitude of multiple parasitoids on host suppression for each heterospecific parasitoids assemblage* (parasitoid abbreviations: A: *Asobara sp.*, L: *Leptopilina sp.*, G: *Ganaspis sp.*) and host species (*D. birchii*, *D. pallidifrons*, *D. simulans.).* Green corresponds to observed effect magnitude with multiple parasitoid species, while red corresponds to estimated effect magnitude of multiple parasitoid species. Observed values smaller than estimated translate to risk enhancement for the host with multiple parasitoid species, while observed values bigger than estimated reflect risk reduction. Big dots represent the means (±SE), and small dots represent raw data. Significant differences between observed and estimated values were calculated with matched-pairs t tests (P-values: *** < 0.001, ** < 0.01, * < 0.05, ns > 0.5).

**Table S1.** *Effects of community structure on the host degree of infestation in the community module model* with the effects of host and parasitoid treatments (two levels each) and their interaction. Abbreviations: 1P: single parasitoid species, 2P: two parasitoids heterospecific, 1H: single host species, 2H: alternative host species.

| **Contrast** | Odds Ratio | p-value |
|---|---|---|
| 2P 1H / 1P 1H | 1.58 | 0.076 |
| 1P 2H / 1P 1H | 0.96 | 0.984 |
| 2P 2H / 1P 1H | 1.32 | 0.376 |

**Table S2.** *Effects of community structure and host species identity on the host degree of infestation.* Effects are shown by the summary of Likelihood-ratio chi-square tests on the species-specific community module model with the effects of host and parasitoid treatments (two levels each), host species (three levels), and their interaction. Degrees of freedom (Df) are given for each factor and the residuals.

| **Effects** | **$\chi^2$** | **Df** | **p-value** |
|---|---|---|---|
| Host treatment | 0.56 | 1 | 0.455 |
| Parasitoid treatment | 7.13 | 1 | 0.008 |
| Host species | 0.07 | 2 | 0.965 |
| Host:Parasitoid | 0.21 | 1 | 0.644 |
| | | 420 | |

**Table S3.** *Effects of community composition on the host degree of infestation.* Effects are shown by the summary of Likelihood-ratio chi-square tests on the community composition model with the effects of host and parasitoid species assemblages (six levels each), and their interaction. Degrees of freedom (Df) are given for each factor and the residuals.

| **Effects** | **$\chi^2$** | **Df** | **p-value** |
|---|---|---|---|
| Host assemblage | 6.21 | 5 | 0.286 |
| Parasitoid assemblage | 32.70 | 5 | < 0.0001 |
| Host:Parasitoid | 68.21 | 25 | < 0.0001 |
| | | 390 | |

**Supporting Text S2:** *Effects of community structure and composition on host suppression*

Probability of host infestation responded differently depending on parasitoid species assemblage (community composition model: $\chi 2(5) = 32.70$, $P < 0.0001$), and the interaction between parasitoid and host assemblages ($\chi 2(25) = 68.21$, $P < 0.0001$).

Neither parasitoid assemblage ($\chi 2(5) = 7.08$, $P = 0.214$), host species assemblage ($\chi 2(2) = 0.58$, $P = 0.748$), nor their interaction ($\chi 2(25) = 4.12$, $P = 0.942$) had a significant effect on *D. birchii* DI (Figure S3a). *Drosophila pallidifrons* DI increased with presence of multiple parasitoids, but only significantly when *Ganaspis sp.* was associated with *Leptopilina sp.* (Post Hoc Odds Ratio (OR) LG/GG = 3.77, P = 0.039, but OR AG/GG = 1.17, P = 0.999; Figure S3b). *Drosophila simulans* DI only significantly increased with presence of multiple parasitoids when either *Leptopilina sp.* or *Asobara sp.* was associated with *Ganaspis sp.* (OR LG/LL = 33.07, P < 0.0001, but OR AL/LL = 4.51, P = 0.105; OR AG/AA = 9.20, P = 0.0007, but OR AL/AA = 0.39, P = 0.318; Figure S3c).

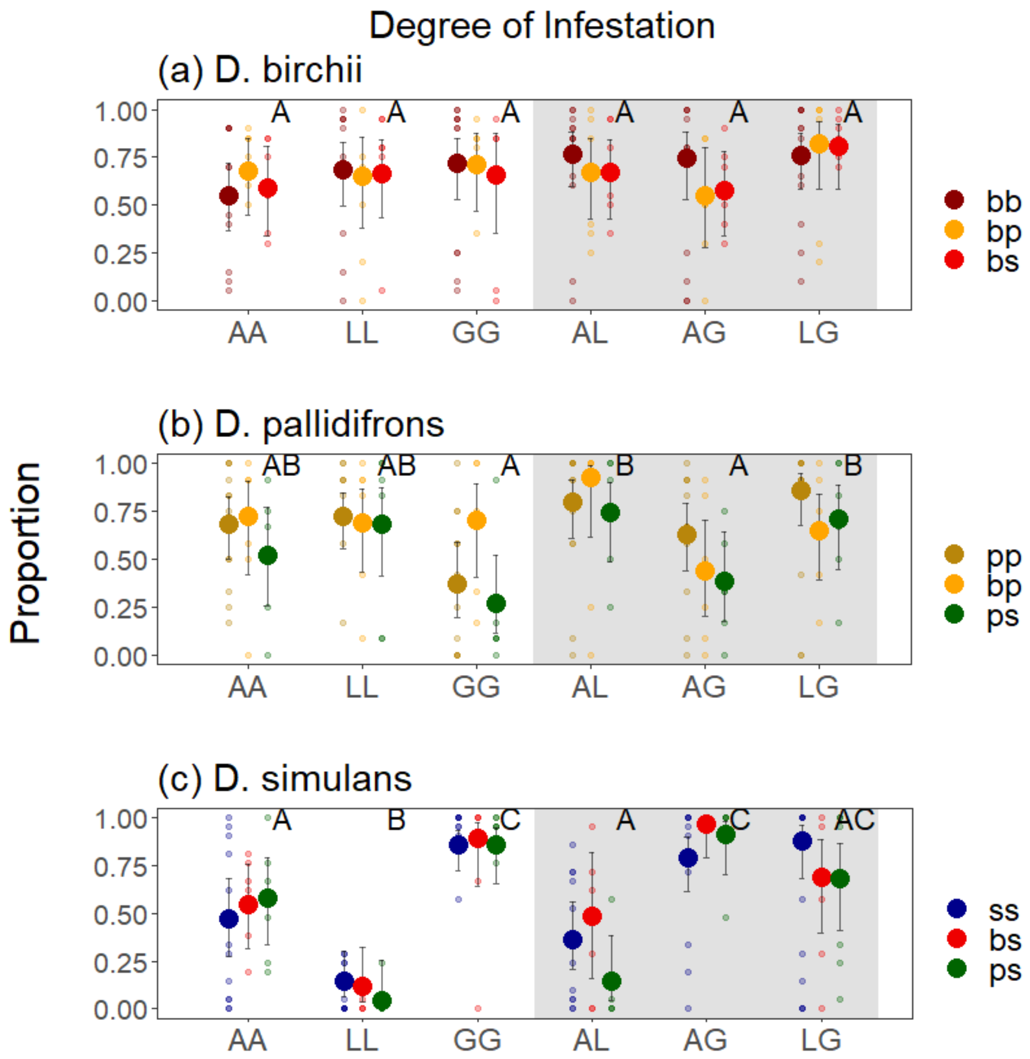

**Figure S3.** *Effects of community composition on the degree of infestation (DI) of each host species* (a) *D. birchii*, b) *D. pallidifrons*, c) *D. simulans*) depending on host assemblage (represented by different colors; host abbreviations: b: *D. birchii*, p: *D. pallidifrons*, s: *D. simulans*) and parasitoid assemblage (on the x axis; parasitoid abbreviations: A: *Asobara sp*., L: *Leptopilina sp*., G: *Ganaspis sp*.). Different capital letters denote significant differences between parasitoid assemblage (from community composition models). White/grey panel: without/with interspecific exploitative competition between parasitoid species. The small points represent the observed values, and the large points represent the predicted values with their 95% confidence intervals

**Table S4.** *Effects of community structure on the probability of successful parasitism.* Effects are shown by the summary of Likelihood-ratio chi-square tests on the community module model with the effects of host and parasitoid treatments (two levels each) and their interaction. Degrees of freedom (Df) are given for each factor and the residuals.

| **Effects** | $\chi^2$ | **Df** | **p-value** |
|---|---|---|---|
| Host treatment | 3.40 | 1 | 0.065 |
| Parasitoid treatment | 0.22 | 1 | 0.635 |
| Host:Parasitoid | 1.614 | 1 | 0.204 |
| | | 637 | |

**Table S5.** *Effects of community structure and host species identity on the host degree of infestation.* Effects are shown by the summary of Likelihood-ratio chi-square tests on the species-specific community module model with the effects of host and parasitoid treatments (two levels each), host-parasitoid pairs (nine levels), and their interaction. Degrees of freedom (Df) are given for each factor and the residuals.

| **Effects** | $\chi^2$ | **Df** | **p-value** |
|---|---|---|---|
| Host treatment | 0.00 | 1 | 1 |
| Parasitoid treatment | 0.00 | 1 | 1 |
| Host-parasitoid (HP) | 85.33 | 8 | < 0.0001 |
| Host treat:Parasitoid treat | 0.00 | 1 | 1 |
| Host treat:HP | 24.01 | 8 | 0.002 |
| Para treat:HP | 16.84 | 8 | 0.032 |
| Host treat:Para treat:HP | 36.81 | 8 | < 0.0001 |
| | | 606 | |

**Supporting Text S3:** *Effects of community structure and composition on parasitoid performance*

Successful parasitism rates significantly increased in modules with a parasitoid competitor for *Asobara sp*. on *D. simulans* with the presence of *Leptopilina sp*. (Post Hoc Odds Ratio (OR) = 4.53, P = 0.003) and *Ganaspis sp*. (OR = 3.88, P = 0.008) in the exploitative competition modules compared to the host-parasitoid pair in isolation, but not in modules with both exploitative competition and alternative host (Figure 4c). The increase in successful parasitism rate of *Asobara sp*. on *D. simulans* in the exploitative competition modules could be due to the suppression of the *D. simulans* immune response by *Leptopilina sp*. and *Ganaspis sp*., and the contrasting results when an alternative host was present could be due to differences in oviposition behavior (some parasitoid randomly lay eggs in many hosts while Braconid species are more specialized in certain groups of hosts), although those mechanisms were not tested in the present study.

Successful parasitism rates of *Ganaspis sp*. on *D. simulans* also significantly increased in modules with a parasitoid competitor compared to the host-parasitoid pair in isolation, but only in both exploitative competition and alternative host modules, and for certain species assemblages. It increased with the presence of *Leptopilina sp*. only when *D. simulans* was associated with *D. birchii* (OR = 13.69, P = 0.004), and marginally with the presence of *Asobara sp*. only when *D. simulans* was associated with *D. pallidifrons* (OR = 4.70, P = 0.053) (Figure 4i).

Successful parasitism rates of *Leptopilina sp*. on *D. birchii* significantly increased with presence of *Asobara sp*., but only in modules with both exploitative competition and alternative host (when *D. birchii* was associated with *D. pallidifrons* OR = 4.83, P = 0.0001, and when *D. birchii* was associated with *D. simulans* OR = 3.86, P = 0.0003), and was not significantly affected by the presence of *Ganaspis sp*. for any of the host combinations (Figure 4d).

Successful parasitism rates of *Ganaspis sp*. on *D. birchii* was significantly affected by the presence of *Asobara sp*., but only in modules with both exploitative competition and alternative host, and the direction of the effect depended on the alternative host species. It increased when *D. birchii* was associated with *D. pallidifrons* (OR = 4.69, P = 0.004), but decreased with presence of *Asobara sp*., when *D. birchii* was associated with *D. simulans* (OR = 0.13, P = 0.016), suggesting antagonistic interactions between those species with that host assemblage (Figure 4g).

Successful parasitism rates of the other eight host-parasitoid pair did not significantly differ with either or both the presence of a parasitoid heterospecific (exploitative competition modules) and of an alternative host (alternative host modules) compared to the module with the host-parasitoid pair in isolation (Figure 4).
